# Pigment-Dispersing Factor is present in circadian clock neurons of pea aphids and may mediate photoperiodic signalling to insulin-producing cells

**DOI:** 10.1101/2023.04.27.538577

**Authors:** Francesca Sara Colizzi, Jan A. Veenstra, Gustavo L. Rezende, Charlotte Helfrich-Förster, David Martínez-Torres

**Author notes:** **Corresponding authors:** (Charlotte Helfrich-Förster) (David Martinez-Torres).

## Abstract

The neuropeptide Pigment-Dispersing Factor (PDF) plays a pivotal role in the circadian clock of most Ecdysozoa and is additionally involved in the timing of seasonal responses of several photoperiodic species. The pea aphid, *Acyrthosiphon pisum,* is a paradigmatic photoperiodic species with an annual life cycle tightly coupled to the seasonal changes in day length. Nevertheless, PDF could not be identified in *A. pisum* so far. In the present study, we identified a PDF-coding gene that has undergone significant changes in the otherwise highly conserved insect C-terminal amino acid sequence. A newly generated aphid-specific PDF antibody stained four neurons in each hemisphere of the aphid brain that co-express the clock protein Period and have projections to the *pars lateralis* that are highly plastic and change their appearance in a daily and seasonal manner, resembling those of the fruit fly PDF neurons. Most intriguingly, the PDF terminals overlap with dendrites of the insulin-like peptide (ILP) positive neurosecretory cells in the *pars intercerebralis* and with putative terminals of Cryptochrome (CRY) positive clock neurons. Since ILP has been previously shown to be crucial for seasonal adaptations and CRY might serve as a circadian photoreceptor vital for measuring day length, our results suggest that PDF plays a critical role in aphid seasonal timing.

## 1. Introduction

The neuropeptide Pigment-Dispersing Factor (PDF) plays a pivotal role in the circadian clock of most insects investigated so far [1–4]. In fruit flies and cockroaches, PDF is an essential output molecule of specific circadian clock neurons that control behavioural rhythmicity and serves additionally as a communication factor within the circadian clock network [3,5–7]. Furthermore, in several insects, including fruit flies, PDF is implicated in seasonal timing (photoperiodicity) [8–14] and, in one moth species, in annual rhythms [15]. Besides that, PDF is involved in negative geotaxis [16], long-term memory [17,18], pheromone production [19], renal physiology [20] and general metabolism [21] of fruit flies.

Photoperiodism is the ability to perceive day length (photoperiod) as an anticipatory cue of seasonal changes and to respond with appropriate physiological and behavioural adjustments [22]. The ability to measure day length is essential for survival in temperate regions of the planet. Measuring day length requires an endogenous timing system as a reference, and the circadian clock is hypothesized to fulfil this role [23]. In the blow fly *Protophormia terraenovae*, the ablation of the PDF-positive clock neurons renders the flies unable to discriminate between long and short days [24] and thus unable to prepare in time for the coming winter. In the bugs *Plautia stali* [12] and *Pyrrhocoris apterus* [13], PDF is essential for entering overwintering diapause under short days, while in female mosquitoes (*Culex pipiens*) [25] and fruit flies [11], PDF is necessary to maintain the ability to reproduce during long summer days. Fruit flies and cockroaches (*Rhyparobia maderae*) have higher PDF levels/ denser PDF branching patterns under long summer days than under short winter days [14,26]. All these findings suggest that PDF conveys day length information to the photoperiodic system: in some insects PDF appears to signal long days, while in others it rather signals short days.

Aphids (Hemiptera: Aphididae) were the first animals described as photoperiodic [27]. In temperate regions of the planet, they adapt to seasonal changes with a remarkable life cycle. During spring and summer, characterized by longer days, aphid populations consist exclusively of viviparous females that reproduce parthenogenetically (i.e. they are clonal). New embryos are already developing inside the yet unborn older embryos developing inside parthenogenetic aphids, which ensures rapid and efficient reproduction. When autumn comes and the days shorten, aphids switch their reproductive mode and generate males and oviparous sexual females. Adult males and females mate and produce cold-resistant eggs that overwinter and survive the unfavourable season. In the next spring, those eggs hatch and the newly born nymphs initiate a new series of viviparous parthenogenetic female generations that succeed one another until the next autumn.

Despite the remarkable ability of aphids to measure day length, they seemed to lack PDF. Neither the *pdf* gene [28] nor the PDF peptide [29] have been detected in aphids to date, although its putative receptor was identified [30,31]. We hypothesized that the lack of this neuropeptide could be a possible cause for the weak activity rhythms of aphids that dampen quickly after transfer into constant darkness [32]. However, other species of the order hemiptera do have PDF. For example, the cicada *Meimuna opalifera* [33] and different species of bugs, such as *Gerris paludum*, *Rhodnius prolixus*, and *Riptortus pedestris* [34–36] have PDF. Some of these species are strongly photoperiodic, and as mentioned above, PDF appears to be involved in their photoperiodic responses. These findings suggest that also aphids may possess PDF but that its coding sequences may have diverged during evolution so much that neither a similarity-based BLAST search, nor antibodies recognizing PDFs in other insects were able to identify aphid PDF. This happened in the beetle *Tribolium castaneum*, for which initially no PDF gene sequence could be identified [37], while, as in aphids, the G protein coupled PDF receptor was immediately detected [38]. Finally, a comparative analysis of several beetle genome sequences and RNAseq assemblies identified the predicted PDF sequence in *T. castaneum* and revealed that it had undergone significant changes, especially in the C-terminal amino acid sequence [38]. Indeed, when amino acid sequences are poorly conserved, identifying orthologous short neuropeptides may become very difficult. It is therefore not surprising that the *Acyrthosiphon pisum pdf* gene was initially not recognized as such, even though its presence was strongly suggested by the existence of a gene encoding a *pdf* receptor in its genome [28]. However, thanks to the current availability of genome and transcriptome assemblies for diverse aphid and aphid-related species, we report here the finding of a highly divergent *pdf* gene in the pea aphid genome that was already present in the ancestor of all the Aphidomorpha.

Using a newly generated antibody against the predicted amino acid sequence of *A. pisum* precursor PDF, we were able to stain four neurons in each lateral protocerebrum of the aphid brain that co-express the clock protein Period and show projections into the superior protocerebrum, strongly resembling those of the fruit fly PDF neurons. They overlap with fibres of Cryptochrome positive clock neurons in the lateral and superior protocerebrum and with the dendrites from the insulin-like peptide positive neurosecretory cells in the *pars intercerebralis*. The PDF terminals in the superior protocerebrum are highly plastic and change their length in a daily and seasonal manner. Together, our results establish the presence of PDF in aphids and suggest that PDF plays a pivotal role in the circadian and seasonal timing of pea aphids.

## 2. Results

### 2.1 Pea aphids possess a highly divergent PDF encoding gene

As typical for neuropeptides, PDF is synthesized from a larger, inactive precursor protein (prepro-PDF), which consists of a signal peptide and a PDF-associated peptide (PAP) followed by the region that, after processing, will constitute the mature PDF [40]. The signal peptide guides the protein to the secretory pathway and is later cleaved off. Similarly, the mature PDF is cut out from the PAP by neuropeptide convertases that recognise single or paired basic residues as cleavage sites [41] rendering a peptide consisting of 18 amino acids highly conserved across Arthropoda [40]. Finally, the neuropeptide is further processed and amidated at its C-terminal end to become biologically active [42].

Our Tblastn searches for pea aphid PDF on the genome assembly yielded a small protein sequence that had rather limited sequence similarity to PDF. Using this sequence as a query in a Tblastn search on transcriptome shotgun assemblies of Aphidomorpha identified a number of aphid transcripts. The putative aphid PDF sequences in these transcripts are conserved and they all start with a signal peptide. This strongly suggested that these are the aphid PDF precursors. Our BlastP searches with theses putative PDF precursors using the NCBI refseq_protein database restricted to *A. pisum* (AL4 genome assembly [39]) (see Materials and Methods), yielded two hits. These corresponded to two predicted proteins (XP_003244595.1 and XP_016659923.1) described as “uncharacterized protein LOC100574816” isoforms X1 and X2 respectively of which the first one is an ortholog of the putative PDF precursors (Fig. 1). The N-terminal convertase Lys-Lys-Lys cleavage site flanking the sequence of the putative mature PDF in the predicted proteins (see Fig.1) is very unusual [41], while the internal Arg-Arg cleavage site, if cleaved, would yield a very short PDF (11 amino acids) or, if left intact, the C-terminal end of the molecule would be very different from other insects (see Fig. 1). In earlier attempts to identify this gene, it appeared that the partial sequence similarity with PDF was fortuitous rather than reflecting a genuine PDF neuropeptide, since neither of these possibilities seemed to yield a credible PDF neuropeptide. The presence of orthologous transcripts in other aphid species, and even the more distantly related grape phylloxera *Daktulosphaira vitifoliae*, showed the putative neuropeptide to be conserved and, hence, likely functional.

**Figure 1:**
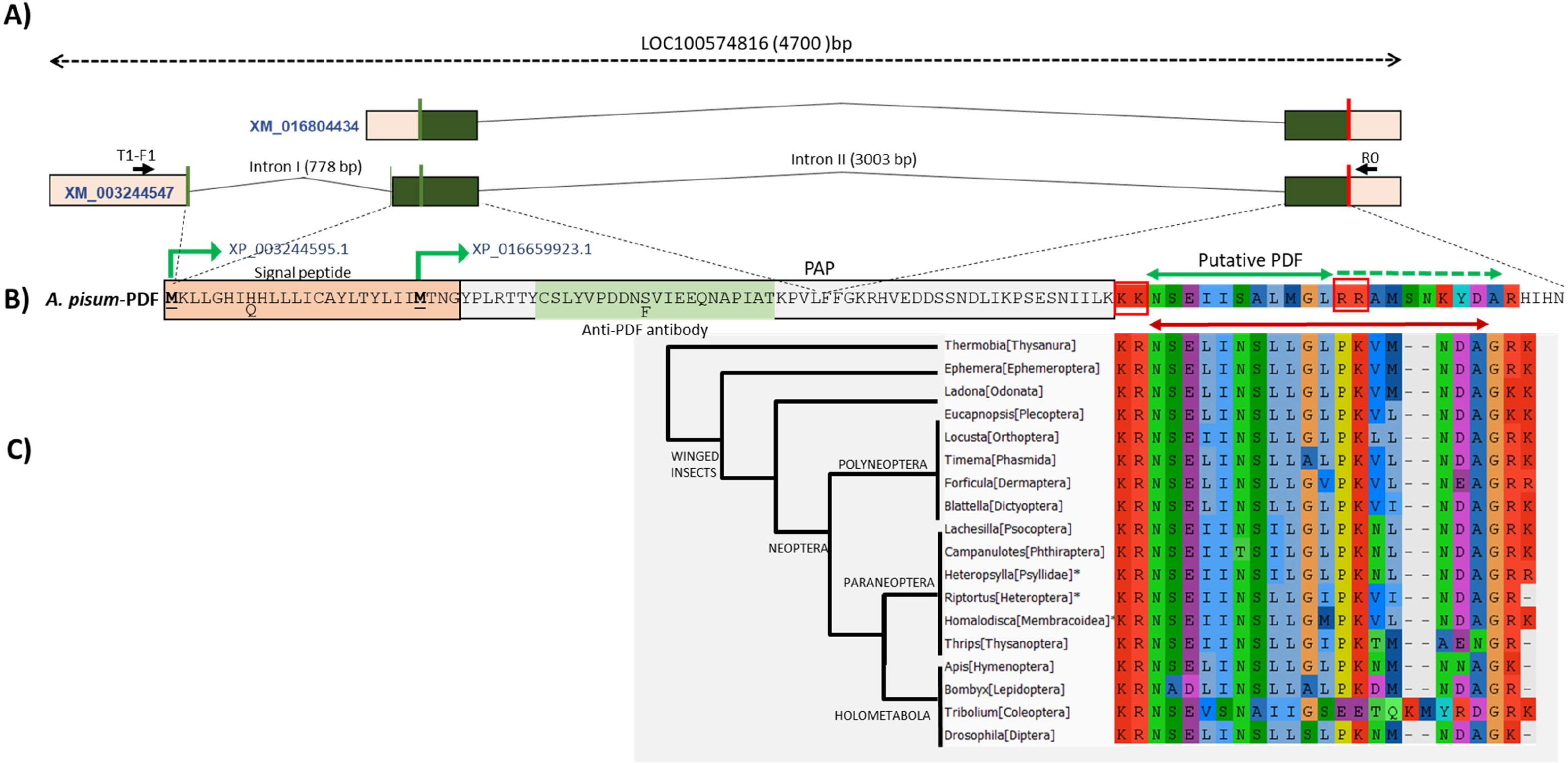
Identification and characterisation of PDF in *A. pisum*. **A:** Schematic representation of the two predicted transcripts (indicated by accession numbers in blue) that encode predicted proteins that partially aligned with query sequences in BlastP searches. Predicted exons are indicated in coloured boxes. Dark green in exons correspond to CDSs. Vertical green lines correspond to initial methionines and vertical red lines to stop codons. Black lines connecting exons indicate predicted introns (size indicated in parenthesis). The position of the primers used to experimentally characterise the transcripts are indicated by black arrows. **B:** Predicted protein isoforms (indicated by their accession numbers in blue) from the two predicted transcripts in A. Both proteins start at different methionines (green angled arrows). Dotted lines connect regions of the protein encoded by CDSs in particular exons. Different elements of the PDF precursor are indicated: signal peptide, PAP (PDF Associated Propeptide). The carboxyl-terminal region that partially aligns with insect PDF appears in background colours. A putative mature PDF peptide (see sections 2.1, 2.2 and Fig. S1) is indicated by a double green arrow limited by red boxes that correspond to predicted convertase cleavage sites. Dotted green arrow indicates a possible extension of a hypothetical mature peptide if the second convertase cleavage site is not cleaved. Two polymorphisms found in different strains are indicated below the main predicted sequence. The green shadowed sequence indicates the peptide used to raise the pea aphid specific antiserum. **C:** PDF sequences, along with flanking basic residues, from 18 insect species representative of major insect orders, aligned with the *A. pisum* predicted peptide. The double red arrow indicates the extension of the mature PDF in these insects (note the highly divergent sequence of *Tribolium*). A manually built cladogram besides the alignment shows the relationships between insect groups according to [45] (only the genus and order are indicated). * indicates three representatives from Hemiptera. Close to some branches, in capital letters, main insect groupings are indicated. NCBI accession numbers for the included sequences are, from top to bottom: JT495639, GAUK02023238, KAG8239166, GIEC01052504, AKN21252, CAD7590987, GAYQ02044840, PSN52637, GCWJ01020925, GCWD01026746, GCXB01024081, BAN82692, XP_046677079, XP_034243662, XP_006570344, NP_001036920, EFA10486, O96690.

The two pea aphid predicted proteins correspond to conceptual translations from two alternative predicted transcript variants of gene LOC100574816 that spans over 4707 bp (Fig. 1). Predicted transcript XM_003244547 spans 926 nucleotides (nts) distributed in three exons (Fig.1A). This mRNA is predicted to contain a 339 nts coding sequence (CDS) who’s initial ATG codon is distributed between the end of the first exon (AT) and the beginning of the second one (G) (see Fig. 1A and B). Thus, this mRNA can be translated into predicted protein XP_003244595.1 consisting of 112 amino acids (see Fig. 1B). Adding further evidence to this gene encoding a true pea aphid PDF, this protein contains the signal peptide (as predicted by SignalP 6.0 [43]) typical of PDF precursor proteins, apart from the two protein convertase cleavage sites discussed above, necessary to get the mature PDF from the propeptide (see Fig. 1B).

The second predicted transcript (XM_016804434) would yield the predicted protein isoform XP_016659923.1, which is identical in all its length to protein XP_003244595.1 but lacking most of the predicted signal peptide (see Fig. 1A). Given its questionable functionality, it might likely correspond to an artefactual prediction and thus, we excluded it from further analysis.

To experimentally validate the *A. pisum* putative PDF transcript predictions, we designed specific primers based on the 5’ and 3’ UTRs of the predicted transcript XM_003244547 (see Fig. 1A) and used them to PCR amplify the corresponding transcripts from cDNAs synthesised using total RNA purified from different pea aphid strains (see Materials and Methods). Our primer combination yielded amplified fragments of the expected size according to the prediction.

We performed direct sequencing of the PCR-amplified fragments from the main transcript (XM_003244547), that encodes the full putative pre-propeptide, from seven pea aphid strains from diverse geographical origins (see Materials and Methods), including the LSR1 strain, whose genome was firstly published [44]. All sequences were deposited in GeneBank (Accession numbers indicated in Table S1). For all the strains, our experimental sequences coincided, for the most part, with the predicted sequence. However, we found two nonsynonymous polymorphisms at amino acid positions 8 and 43, that, in the predicted transcript correspond to histidine (H) and serine (S) respectively (Fig. 1B). We found that five out of the seven strains sequenced (including the reference strain LSR1) were in fact heterozygous at both positions having, in addition to the predicted H and S, glutamine (Q) and phenylalanine (F) respectively at those two positions (see Fig. 1B). Sequencing of the cloned sequences revealed that alleles H and S reside on the same chromosome, and Q and F on the other one. Coincident with these results, two strains were homozygous at both positions. Strain BOL was QF while strain GR was HS (Table S1). It is yet unclear the relevance, if any, of these polymorphisms (but see below).

Finally, our experimental sequences from the seven pea aphid strains perfectly matched the 3’ end sequence of transcript predictions that correspond to the carboxyl-terminal end of the predicted PDF propeptide containing the putative mature PDF described above. Figure 1C shows the alignment of this region in the pea aphid with 18 sequences representative of major insect orders. This region in the pea aphid seems to have diverged much when compared with other insect groups (with the exception of *Tribolium*, which also possesses a rather divergent PDF, see Fig. 1C) especially after the second putative convertase cleavage site. Indeed, for the 25 amino acid positions aligned in Fig. 1C, the average number of differences among the 17 insect species (excluding both *Tribolium* and *A. pisum*) is 4.3 (ranging from 1 to 9). However, the average number of differences between the *A. pisum* sequence and the rest is 13.1 (ranging from 12 to 15).

### 2.2 A divergent PDF is characteristic of Aphidomorpha

To investigate whether the putative PDF found in Blast searches in the pea aphid was also present in other aphid species we performed BlastP or tBlastn searches in different aphid databases using as query the pea aphid sequence identified in the above section (see Materials and Methods). We found highly similar sequences to the putative *A. pisum* PDF in all searched aphid databases including different species of the two tribes in the subfamily Aphidinae (i.e. Macrosiphini, to which *A. pisum* belongs, and Aphidini), representatives of Eriosomatinae and Lachninae (two distantly related subfamilies within the Aphididae [46]), and, most relevant, in representatives of oviparous families Adelgidae and Phylloxeridae, which separated from true aphids some 200 million years ago [47] (see Fig. S1). These results led us to conclude that we had indeed found the pea aphid PDF. Furthermore, the 11 amino acids, flanked by the two convertase cleavage sites, present in the *A. pisum* sequence, are identical in most aphid sequences including the Adelgidae and Phylloxeridae representatives (see Fig. 1C and S1). Interestingly, the unusual Lys-Lys cleavage site observed in the *A. pisum* sequence is also present in most other aphid species, but is replaced by conventional Lys-Arg in oviparous families Adelgidae and in the grape phylloxera *Daktulosphaira vitifoliae* and also in the single representative of the distantly related aphid subfamily Eriosomatinae (see Fig. 1C and Fig. S1). This degree of conservation in the sequence delimited by the two putative convertase sites points to this short peptide being the aphid PDF active form, although additional experiments should confirm this hypothesis

Thus, although the PDF neuropeptide is different from that of other insect species, within the Aphidomorpha it is well conserved. However, the remainder of the PDF precursor has evolved significantly in aphids. In fact, a phylogenetic tree built using the whole PDF precursor sequences (Figure S1), recovers the main aphid groups and known evolutionary relationships among them. As expected, the most divergent sequences correspond to representatives of basal families Adelgidae and Phylloxeridae, in the latter case to the point that an unambiguous signal peptide is no longer predicted by SignalP 6.0. Thus, our results point to a divergent PDF neuropeptide already present in the ancestor of all Aphidomorpha and the gene evolving in the group since then.

### 2.3 The PDF antibody labels four neurons in the lateral brain that slightly differ in size

Using the newly generated antibody against 21 amino acids of the *A. pisum* PDF precursor (see Fig. 1B) on brain whole-mounts, we found four PDF-immunoreactive (PDF-ir) cell bodies per hemisphere, located between the central brain and the optic lobe (Fig. 2A, B). These neurons slightly differed in size. Two of the four PDF-ir neurons possessed large cell bodies (area of 49 to 70 µm^2^, mean size 57.4 µm^2^ at the confocal plane showing the maximal size of the cell), the other two were clearly smaller (26 to 47 µm^2^, mean size 37 µm^2^) (Fig. 2B). Sometimes the two smaller cells had somata of similar size, but often we could distinguish one of intermediate size and one of a rather small size (Fig. 2B’, B’’). The somata of the four cells were always very close together and the neurites originating from them largely intermingled with each other, so that it was impossible to follow the neurites from individual neurons. Nevertheless, we were able to count the number of fibres in certain fibre tracts. The neurites of all four neurons appeared to innervate a neuropil located in front of the lobula complex and proximally to the medulla that strongly resembled the accessory medulla (AME) of other insects [48–50] (Fig 2 A, B, D, E). In the following, we will call this structure AME-like region. We never saw any neurites running beyond the AME-like structure and entering the optic lobes or the compound eyes (Fig. 2A). At least two neurons from each hemisphere projected to the contralateral AME-like region, respectively (arrow in Fig. 2 A, C, D, E’) and two neurons sent fibres to the *pars lateralis* in the superior protocerebrum, where they terminated by forming prominent varicosities (arrowheads in Fig. 2 A, C-D). The fibres forming the commissure to the contralateral brain hemisphere were clearly distinguishable from the fibres terminating in the *pars lateralis*, because they were in different depths of the brain (Fig. 2C, C’, C’’). Nevertheless, sometimes, a single fibre of the contralaterally projecting neurons followed the ones that terminated in the ipsilateral *pars intercerebralis*, intermingled with their varicose terminals, then left them and followed the fibres running to the contralateral AME-like structure (double arrowhead in Fig. 2E).

**Figure 2:**
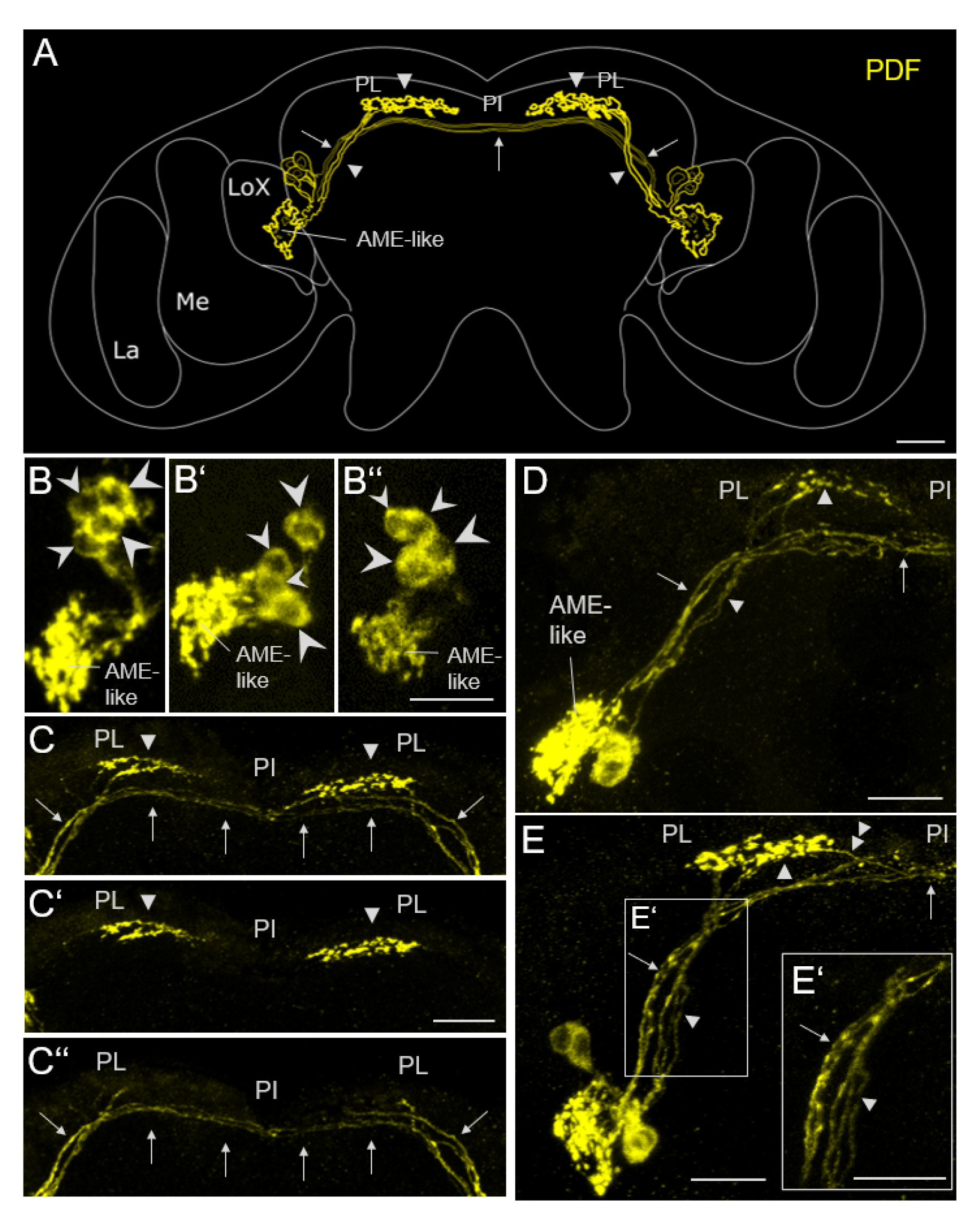
PDF-like immunostaining in the pea aphid brain. A: Schematic representation of the PDF-ir neurons and their neurites. All four neurons appear to send neurites into an accessory medulla-like structure (AME-like). Two main fibre bundles run medially; one crosses in the *pars intercerebralis* (PI) to the contralateral hemisphere and connects both AME-like structures with each other (arrows), the other runs to the *pars lateralis* (PL), where it shows varicose terminals (arrowheads). LA: Lamina. ME: Medulla. LoX: Lobula complex. B: Somata of the PDF-ir neurons from the left brain hemispheres of 3 different aphid brains. Their slightly different soma size is indicated by differently sized arrowheads. C: PDF-ir fibres in the superior protocerebrum. Arrows mark the fibres projecting to the contralateral AME-like and arrowheads mark the fibres ending in the ipsilateral PL. C: Full Z stack (20 stacks, thickness 1.5µm) of the PDF fibres in the superior protocerebrum. C’: overlay of 6 more dorsally located confocal stacks showing the PDF-ir terminals in the PL. C’’ overlay of 7 more ventrally located confocal stacks showing the contralateral PDF-ir fibres. D: left brain hemisphere showing that 2 fibres leaving the PDF-ir somata, run to the PL and terminate there (arrowheads) and 2 others cross via the PI to the contralateral brain hemisphere. E: example of a brain hemisphere, in which 3 fibres run to the PL and one of them continues toward the PI. E’: magnification of the area in the rectangle of E. Scale bars: 20µm.

### 2.4 The PDF-ir cells are Cryptochrome-negative but Period-positive clock neurons

To test whether the PDF-ir cells are clock neurons, we performed co-labelling of anti-PDF with anti-Period (PER) and anti-Cryptochrome (CRY). Aphids were collected in the early morning, one hour before lights-on (ZT23), and in the afternoon, 6 hours before lights-off (ZT11). At these times (especially at ZT11) our previous study has detected significant PER staining in the lateral clock neurons (two CRY-negative LNs and one CRY-positive LN+), in 3 dorsolateral clock neurons (DLNs) and ∼7 dorsal clock neurons (seven CRY-negative DNs and two CRY-positive DN+s) and ∼9 lamina neurons (LaNs) [29]. Besides the one LN+ and the two DN+s, CRY was present in all LaNs (Fig. 3A [29]). Since we did not find PDF in the optic lobes, we did not further consider PER and CRY staining in the lamina in the present study.

**Figure 3:**
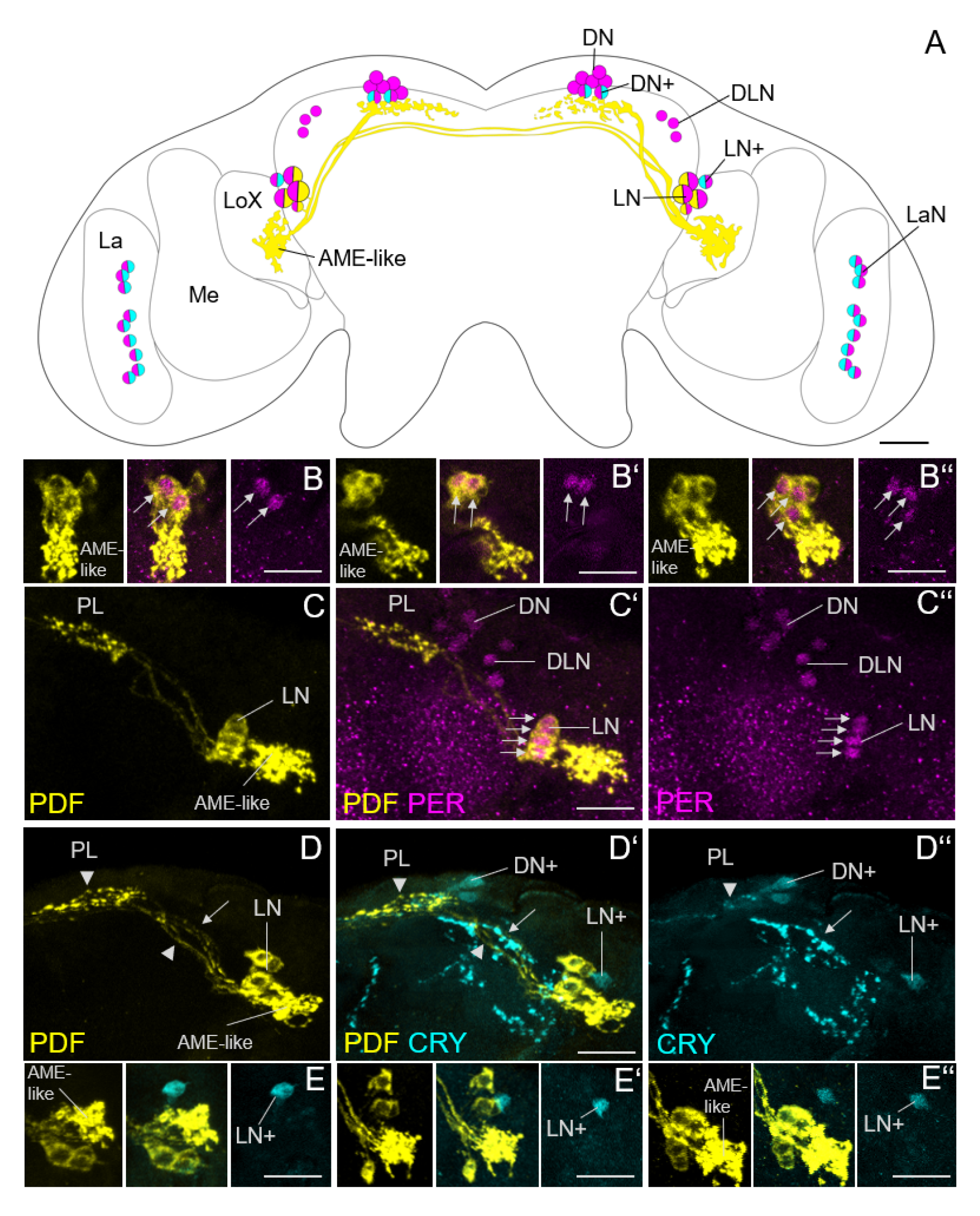
PDF and PER are colocalized in the CRY-negative lateral clock neurons (LN) of the pea aphid brain. A: Schematic representation of the PDF- (yellow), PER-(magenta), and CRY- (cyan) ir neurons. DN: dorsal neurons; DN+ CRY-positive dorsal neurons; DLN: dorsolateral neurons; LN lateral neurons; LN+: CRY-positive lateral neuron; LaN: lamina neurons; La, lamina; Me, medulla; LoX: lobula complex; AME-like: accessory medulla like structure. B-B’’: Overlays of 3 to 5 confocal stacks showing PDF and PER double-labelling in the LNs of three brains stained at ZT23. Arrows mark the double-labelled cells (two LNs in B and B’, and three LNs in B’’. C-C’’: Overlay of 21 confocal stacks showing PDF and PER double-labelling in the right brain hemisphere at ZT11. All four PDF-ir neurons express also PER (arrows). D-D’’: Overlay of 10 confocal stacks depicting PDF and CRY double-labelling in the right brain hemisphere at ZT23. The CRY-positive LN+ is PDF-negative, but the projections of the LN+ overlap with the PDF-positive fibres projecting contralaterally (arrow). In addition, the fibres stemming from the CRY-positive DN+s overlap with the PDF-positive terminals in the *pars lateralis* (PL) (arrowhead). E-E’’: Overlays of 3 to 5 confocal stacks showing PDF and CRY labelling in the LNs and LN+ of three brains. The CRY-positive LN+ was always PDF negative. Scale bars: 20 µm.

Consistent with our previous study, we found PER in the nuclei of the LNs, DLNs and DNs (Fig. 3). At ZT23, PER staining was identical to the previous description and found in the three lateral neurons (two LNs and one LN+). PDF was present in the cytoplasm of the two PER-positive LNs (Fig. 3B). According to the size of the somata, these two PER/PDF positive cells corresponded to the large PDF-ir neurons (Fig. 3B). In rare cases we found a third PER/PDF-positive cell at ZT23 (Fig. 3B’’). At ZT11, PER staining was stronger and usually present in the nuclei of all 4 PDF-positive neurons (Fig. 3C). Co-staining with anti-PDF and anti-CRY showed that PDF and CRY never co-localized in the same neurons, although the neurites of the PDF and CRY-positive cells partly overlapped (Fig. 3D, E).

We conclude that all four PDF-ir neurons are PER-positive clock neurons that don’t express CRY. In our previous study, we had obviously overlooked two of the LNs due to low PER staining intensity.

### 2.5 The PDF-ir fibres overlap with fibres stemming from the CRY-positive clock neurons (LN+ and two DNs)

Although the CRY-positive LN+ was PDF-negative, the fibres arising from it always overlapped with the PDF-positive fibres that projected contralaterally (arrow in Fig. 3D). Furthermore, the fibres stemming from the two CRY-positive DN+s overlapped with the PDF-terminals in the *pars lateralis* (arrowhead in Fig. 3D). This suggests that the CRY-positive and CRY-negative clock neurons communicate with each other.

### 2.6 The PDF-ir terminals in the PL overlap with dendrites from the insulin-like peptide producing neurosecretory cells

Our previous study has shown that insulin-like peptide 4 (ILP4) is a promising candidate for being the predicted virginoparin responsible for the switch between parthenogenesis and sexual reproduction in aphids [51]. The ILP4-producing neurosecretory cells (IPCs) in the *pars intercerebralis* have putative dendritic connections to the *pars lateralis* suggesting a possible communication between the circadian and photoperiodic systems. To elaborate this further, we performed double-immunolabelling with anti-PDF and anti-ILP4 as well as with anti-CRY and anti-ILP4. We found that the putative ILP4 dendrites fully overlap with the PDF terminals in the *pars lateralis* (Fig. 4A) as well as with the CRY-positive fibres arising from the DN+s (Fig. 4B).

**Figure 4:**
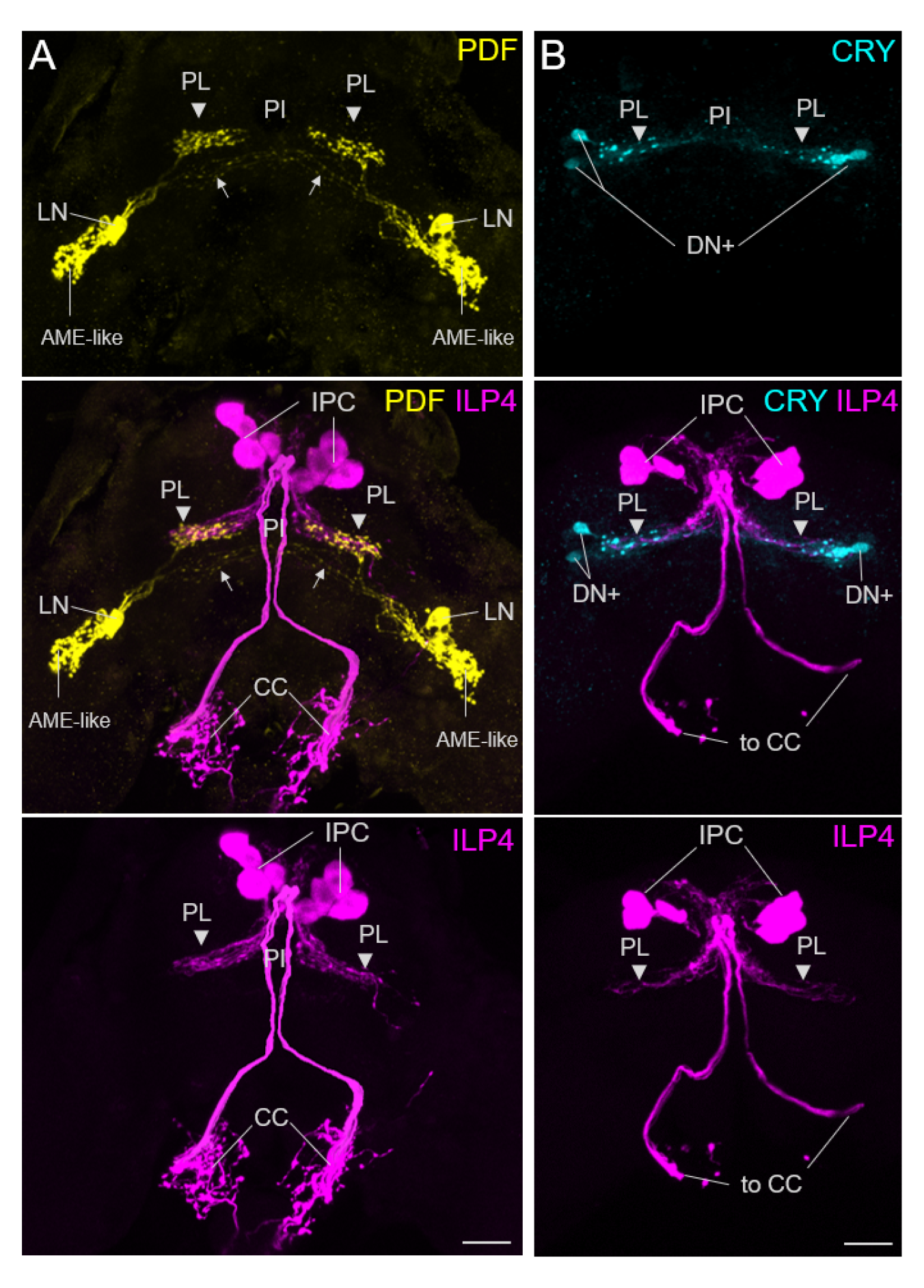
The PDF terminals of the LN and the fibres from the CRY-positive DN+s overlap with putative dendrites of the insulin-like peptide producing cells (IPC) in the *pars lateralis* (PL). A: Overlay of 50 confocal stacks depicting PDF-ir and ILP4-ir in the brain and corpora cardiaca (CC). The ILP4 positive IPCs have dendrites in the PL and send axons to the CC from where they release ILP4 into the circulation. In the PL, their dendrites overlap with the PDF terminals from the PDF-positive LNs (arrowheads). B: Overlay of 30 confocal stacks showing CRY-ir and ILP4-ir in the brain and the projections to the corpora cardiaca (to CC). Only the CRY-positive dorsal clock neurons (DN+) are shown that project to the *pars intercerebralis* (PI) and overlap with the dendrites of the IPCs in the PL. Labelling as in Fig. 3. Scale bars: 20 µm.

### 2.7 Pdf expression shows daily and seasonal differences in abundance

Since PDF has been shown to be involved in daily rhythms as well as in photoperiodism in other insects (see introduction), and pea aphids show diurnal feeding rhythms and are paradigmatic photoperiodic insects, we investigated whether *Pdf* gene expression was affected by the time of day and day length. We compared the expression of the *Pdf* gene in head extracts at four different times of the day in two groups of aphids: aphids reared under long (summer-like) days (LD) and aphids that had been under short days (SD) since they were embryos (see Materials and Methods) (Fig. 5). We found that *Pdf* gene expression was highly dependent on the time of day (ANOVA: F_(3, 16)_ = 7,70; p = 0,002) with the highest expression at ZT16 (on average 2,6 times higher than at other ZTs). Furthermore, *Pdf* expression depended significantly on day length (ANOVA: F_(1, 16)_ = 11,02; p = 0,004). On average, the *Pdf* encoding gene showed double the expression (2,04 times) under SD than under LD (Fig. 5). We conclude that *Pdf* expression shows daily oscillations and that short days strongly induce *Pdf* expression.

**Figure 5:**
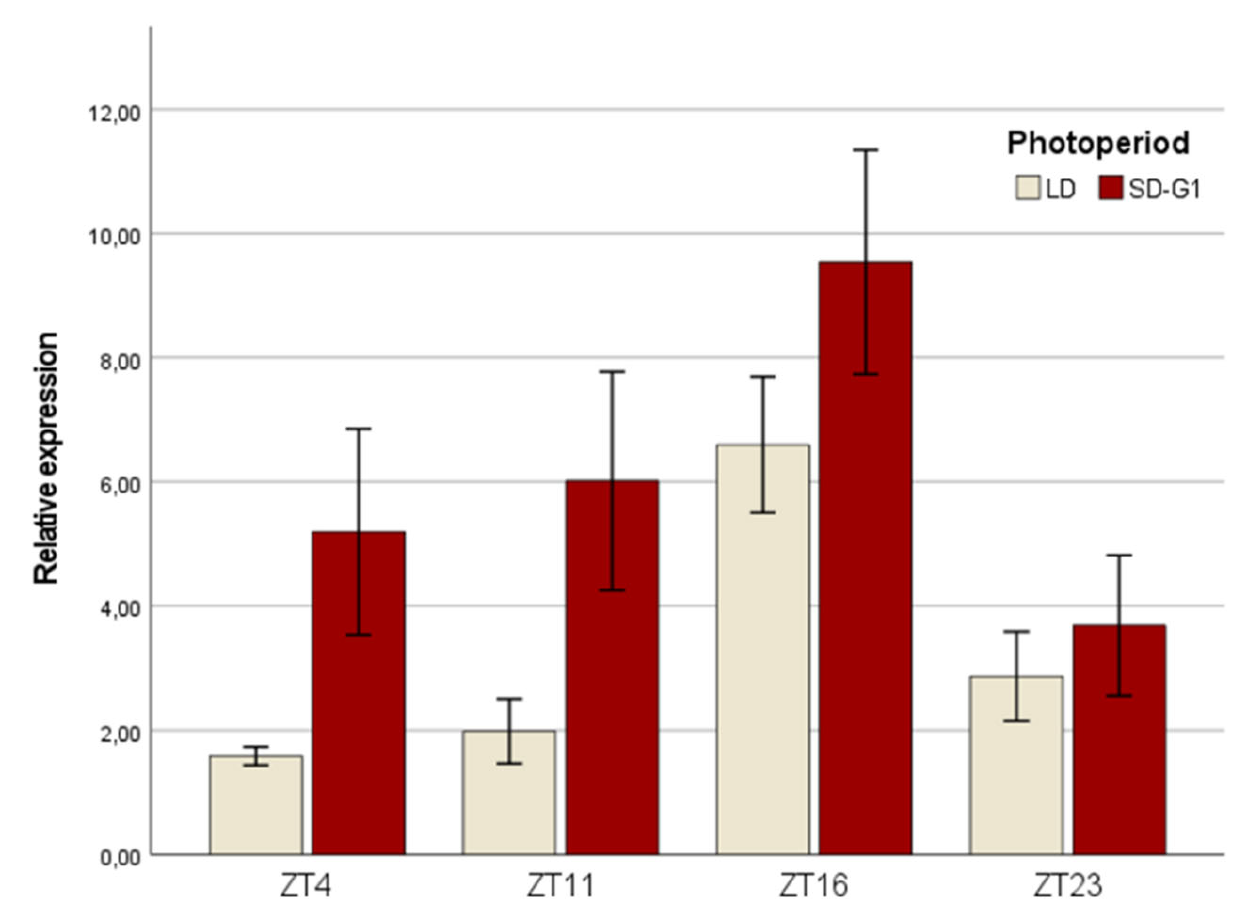
Results from RT-qPCR assays comparing the expression of the PDF coding gene in two photoperiods and 4 timepoints along the day in *A. pisum*. Relative expression levels of the pea aphid PDF coding gene at four timepoints and two photoperiod conditions (see Materials and Methods). Bars are mean values of three replicates +/- SEM.

### 2.8 *The length of the PDF-ir terminals in the* pars lateralis *varies on a daily basis*

After we found that *Pdf* expression shows diurnal oscillations, we aimed to investigate whether the PDF peptide might be used as a circadian clock signal. Therefore, we tested whether the PDF-staining intensity varies during the day and the PDF terminals change their daily shape as was observed in *D. melanogaster* [52,53]. Assuming that PDF peptide abundance peaks after the peak in *Pdf* gene expression, we stained the aphid brains at the expected PDF trough in the early morning, one hour before lights-on (ZT23), and at its expected maximum in the afternoon, 6 hours before lights-off (ZT11). The PDF staining intensity in the PI terminals was determined by three different methods. First, we measured the average pixel intensity in the terminals themselves (see materials and methods) and found no difference between the two time points. However, this method has the disadvantage that it does not consider changes in the shape of the terminals and the visual inspection already showed that the terminals appear wider at ZT23 than at ZT11 (Fig. 6). Therefore, we next used the method of Hermann-Luibl et al. [54], who determined the pixel intensity within a defined area that contained the entire terminals (see Fig. 6A, B). Indeed, we found that the mean staining intensity in this area was significantly higher at ZT23 than at ZT11 (Fig. 6C, left). To confirm this result, we additionally measured the length of the terminals as indicated by ‘double-headed’ arrows in Fig. 6A, B and found that they are significantly longer at ZT23 than at ZT11 (Fig. 6C, right). We conclude that the PDF terminals are plastic and change their shape throughout the day. This makes them suited to transfer daily signals of the lateral clock neurons to downstream neurons.

**Figure 6:**
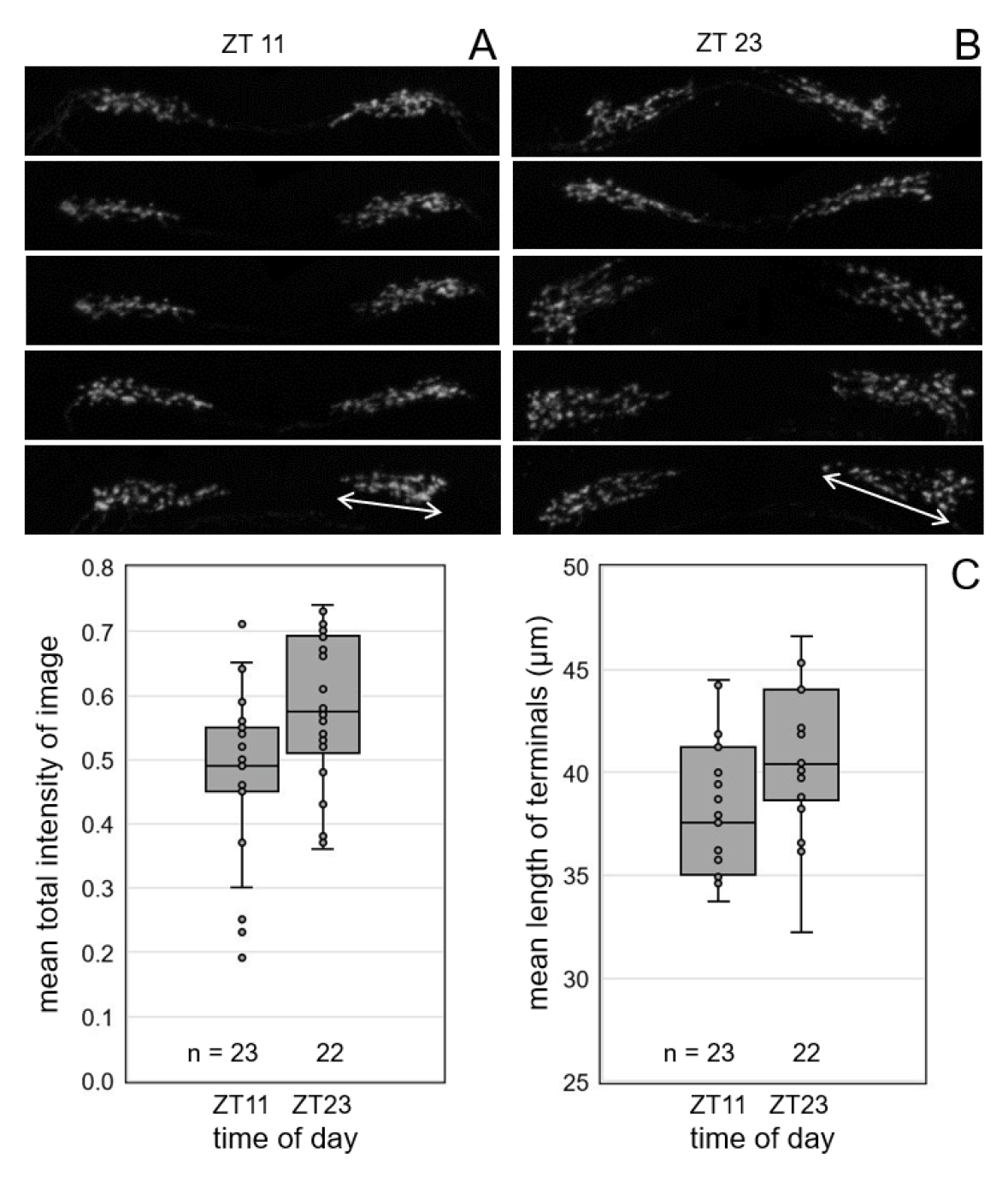
The PDF terminals in the *pars lateralis* are plastic and change their daily appearance. A, B: appearance of the PDF terminals in five representative brains at Zeitgeber Time (ZT) 11 and ZT 23, respectively. mEach image represents an overlay of 10 to 15 confocal stacks. C: mean staining intensity of all images containing the PDF terminals (left) and mean length of the terminals (right) at ZT11 and ZT23, respectively. The number (n) of quantified brains is indicated below the boxplots. The differences in intensity and terminal length were significant (two-sample t-test and Kruskal-Wallis test: p = 0.006 and p= 0.01, respectively).

### 2.9 The PDF-ir terminals are longer under short days than under long days

To test, whether the PDF terminals change their size when aphids experience short days, we immunostained the brains of animals maintained under long days and of aphids that were reared under short days. We decided to stain the brains at ZT23, because we had found more prominent PDF terminals at that time. We found that the PDF terminals significantly increased in size under short days as compared to long days (Fig. 7). Under short days, the PDF fibers spread toward the *pars intercerebralis*, and sometimes the fibers stemming from the two brain hemispheres even touched each other (Fig. 7B). This suggests that in aphids, PDF signaling increases under short days when the aphid starts to produce sexual morphs, and that PDF might be the clock factor of aphids communicating day length to the IPCs.

**Figure 7:**
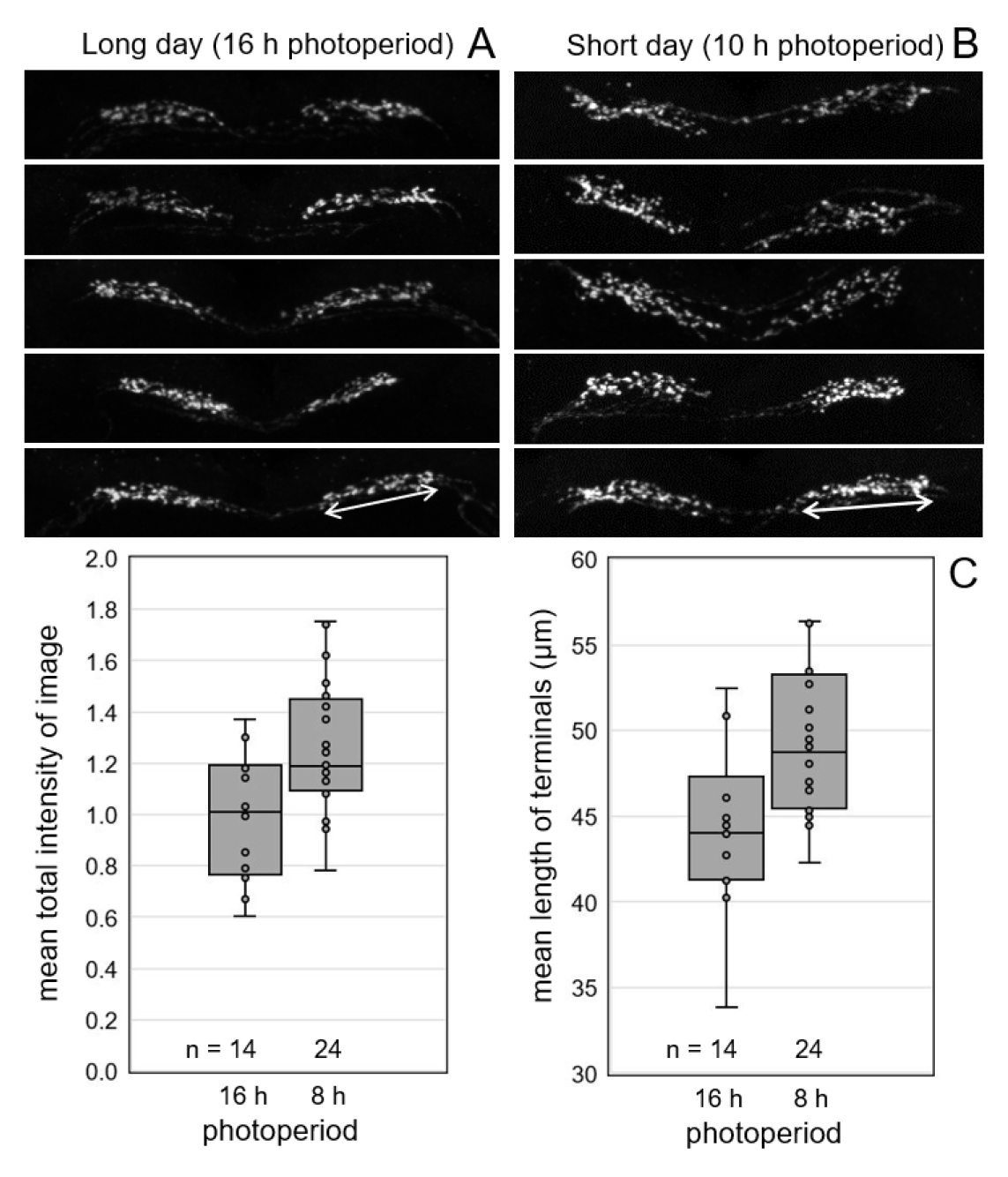
The PDF terminals in the *pars lateralis* are longer under short days than under long days. A, B: appearance of the PDF terminals in five representative brains entrained to long and short photoperiods, respectively. C: mean staining intensity of all images containing the PDF terminals (left) and mean length of the terminals (right) under long and short days, respectively. The number of quantified brains (n) is indicated below the boxplots. The differences in staining intensity and terminal length were significant (two-sample t-test: p = 0.004 and p= 0.004, respectively).

## 3. Discussion

### 3.1 A divergent pdf gene evolved in the Aphidomorpha

The pea aphid, *Acyrthosiphon pisum*, belonged to the few insects in which the *pdf* gene and PDF peptide was not identified. Although the absence of PDF could explain the apparently weak circadian rhythmicity of aphids, it remained questionable why this important peptide, which is present in virtually all panarthropods [3,55–58], should be absent in the strongly photoperiodic aphids. When the *A. pisum* neuropeptide genes were first analysed [28], the PDF gene was not recognized as such because of the significant differences with the other then known insect PDF precursors. A comparative analysis including other aphid and aphid-related species now allowed us to identify its *pdf* gene. The aphid PDF neuropeptide may lack the 7 C-terminal amino acids, that are typical for PDF [42], but its N-terminal sequence is well conserved [58] and its expression pattern in the brain shows large similarities to that of other insect species (see discussion below). There is thus little doubt that this is indeed the aphid PDF gene.

Aphids may be subject to high selective pressures because, as rather static plant suckers, they are particularly exposed to changing environmental conditions and they are special in several aspects. For example, aphids have a rather high number of visual and non-visual pigments that may help them to perceive light, avoid light and be protected from light [59–62]. Most relevant, core clock genes *period* (*per*) and *timeless* (*tim*) experimented high evolutionary rates [63]. Interestingly, these genes participate in the circadian clock feedback loop involved in light-perception. Furthermore, aphids have lost the protein Jetlag, which is involved in the synchronisation of the clock with the daily light-dark cycle (Cortés et al., 2010). However, this rapid evolution is not a general trend of all aphid clock genes since genes involved in the other feedback loop, (i.e. *Clock* and *cycle*) evolved at expected rates [63]. We may speculate that whatever is the selection pressure that drove the divergent evolution of core clock proteins PER and TIM it might have also similarly driven the evolution of the divergent clock neuropeptide PDF. The fact that the levels of *period* gene transcription, similar to the PDF coding gene, are significantly influenced by the photoperiod (with short days inducing higher expression of both genes) speaks for this hypothesis [61,63]. Thus, it is possible that the function of particular clock genes (including PDF) has been directed towards photoperiod-related tasks at the expense of circadian clock ones. Alternatively, relaxed selective constraints may have led to a divergent PDF in the ancestor of all Aphidomorpha (Fig. S1). However, its extreme conservation in all aphid lineages through their ca 200 million years of evolution, points to a strong purifying selection operating to preserve its functionality.

### 3.2 The PDF-positive clock neurons in the aphid brain closely resemble those of other insects

The expression pattern of PDF in aphid circadian clock neurons strongly resembles that of other insects, [64–67]. As true for flies, cockroaches, bugs and bees [12,65,68–70], PDF is present in aphid lateral clock neurons with different soma sizes. These clock neurons send projections to the superior protocerebrum and to the contralateral brain hemisphere, more precisely to a neuropil that strongly resembles the accessory medulla (AME) of other insects. We could not distinguish the projection patterns of the different neurons but it is most likely that those with larger somata project to the contralateral and those with smaller somata remain in the ipsilateral brain hemisphere as was found in cockroaches and flies [49,71]. The AME was first identified as a circadian pacemaker centre in hemimetabolous insects [72–74] and was later established as clock centre in most insects (reviewed in [75]). In these insects, it serves as communication centre for circadian clock neurons and receives photoreceptor input from the compound eyes and extraretinal photoreceptors.

Despite all similarities to other insects, the PDF neurons of aphids are special in the sense that they completely lack PDF fibers in the optic lobes, which would argue in favour of a functional divergence. Furthermore, the AME is particularly rich of varicosities, which are store and release sites of neuropeptides. This speaks against a prominent role of the AME, or at least the PDF fibres in the AME, as a light-input pathway to the circadian clock neurons. Most interestingly, a rather sparse innervation of the optic lobes by PDF fibres was also found in honeybees [50,70]. Furthermore, many varicose endings and no fine dendritic-like fibres were found in the AME of honeybee larvae. This has been interpreted as a lack of photoreceptor input into the honeybee AME, which may be explained by the different lifestyle of bees, which rely much more on social cues than eye-transmitted light-dark cycles to synchronize their circadian clocks [76–78].

Similarly, compound eyes do not appear to play a role as photoreceptors for photoperiodic responses in aphids [79], which may explain the absence of dendritic PDF fibres in AME. Instead, the photoperiodic photoreceptor of aphids was localized to the lateral superior protocerebrum [79], where CRY1-positive clock neurons were later found [29,61]. There, the fibres of these CRY1-positive clock neurons intermingle with fibres of the PDF neurons making it likely that the photosensitive CRY1 synchronizes not only the circadian clock of aphids but at the same time transfers information about day length (photoperiod) to the PDF-positive clock neurons.

Since aphids have a damped circadian clock [32,80], it is possible that PDF plays a weaker role on circadian rhythmicity in aphids compared with other insects. Perhaps the main function of PDF in aphids is the promotion of winter diapause (which in aphids takes the form of sexual reproduction).

### 3.3 PDF as putative factor promoting aphid sexual reproduction in autumn

While both the photoperiodic photoreceptors and the photoperiodic timer appear to localize in the lateral superior protocerebrum (*pars lateralis*) [81], whether the aphid produces sexual or asexual progeny is controlled by neurosecretory cells in the median superior protocerebrum (*pars intercerebralis*) [81]. Ablation of these neurosecretory cells led to the production of sexual morphs even under long days suggesting that they produce a parthenogenesis promoting substance, also called virginoparin [81,82]. Later studies suggested that Insulin-Like Peptides (ILPs) are the virginoparin in question [51,83]. ILP1 and ILP4 are produced in four neurosecretory cells in the *pars intercerebralis* of each brain hemisphere and their expression significantly diminishes under short days promoting sexual reproduction [51]. These neurons project to the corpora cardiaca and from there three nerves (two laterals and one medial) go to the abdomen where ILPs might be released close to the developing aphid embryos, and their levels determine their fate either as parthenogenetic females or as sexual morphs.

Here, we show that the dendrites of the ILP4 expressing neurosecretory cells extend toward the *pars lateralis* where they overlap with the terminals of PDF-positive as well as CRY-positive clock neurons. This strongly suggests that this is the region where the information about photoperiod and time-of day is transferred to these neurosecretory cells. Furthermore, we show that *pdf* expression is significantly higher and that the PDF terminals extend further towards the *pars intercerebralis* under short photoperiods as compared to long photoperiods. Thus, PDF signalling to the ILP neurons may be stronger under short photoperiods and this may in turn result in a decrease of ILP signalling and promote the development of embryos as sexual morphs (Figure 8 summarises this scenario). A similar role of PDF in communicating short photoperiods to the photoperiodic system has been shown for the bugs, *Plautia stali* [12] and *Pyrrhocoris apterus* [13], while very recently, Hidalgo et al. [14] showed that PDF is the long day-signalling factor in *Drosophila melanogaster*. In *D. melanogaster*, the PDF terminals are very prominent under long days, activate the ILP-producing cells in the *pars intercerebralis* and prevent the flies from going into dormancy. Under short days, the PDF terminals become less prominent so that the dormancy inducing factor *Eyes Absent* in the ILP-producing cells can become active and induce dormancy.

**Figure 8:**
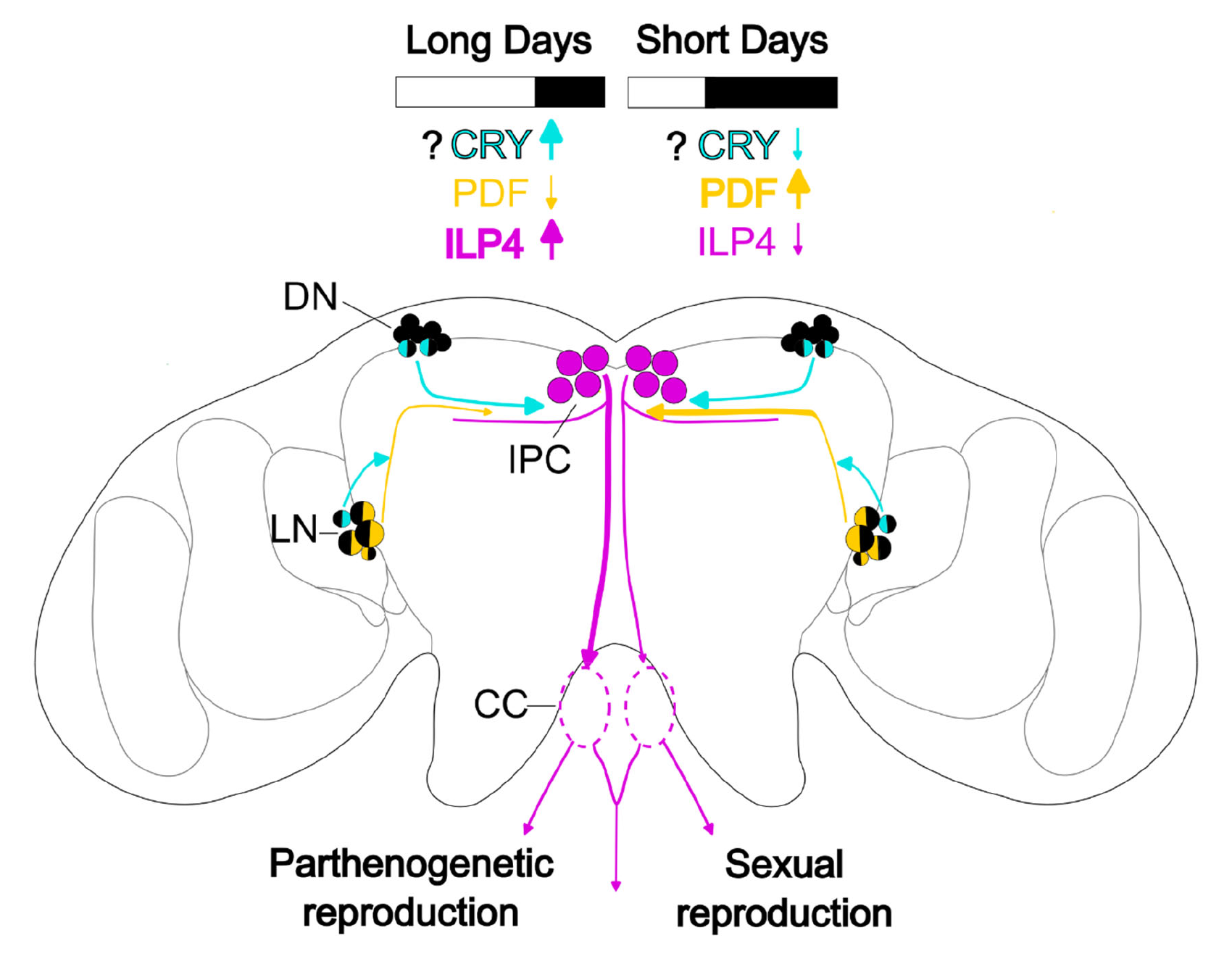
Possible mechanisms of PDF, CRY, and ILP4 signaling during long and short days. On long days (left hemisphere), PDF signaling from the lateral clock neurons (LN) is low, as indicated by the short PDF terminals contacting insulin-producing cells (IPC) in the pars intercerebralis, while ILP4 signaling is high [52]. ILP4 signaling to aphid gonads ensures parthenogenetic reproduction. On short days (right hemisphere), PDF signaling increases, as seen in the long PDF terminals. Consequently, ILP4 signaling is reduced, allowing a switch to sexual reproduction. CRY is expressed in one LN and two dorsal clock (DN) neurons and may function as a seasonal photoreceptor that signals more strongly during long days than short days, but this remains to be experimentally demonstrated.

In summary, we show here that PDF levels could be part of the signal communicating the photoperiod to the *pars intercerebralis* also in aphids. Put in a simple way, PDF may control the synthesis of ILPs to determine the fate of the developing embryos either as parthenogenetic females or as sexual morphs. Further experiments are necessary to prove the function of PDF in the seasonal control of aphid reproduction, but our present results set the stage for future studies into this direction.

## 4. Materials and methods

### 4.1 Aphid strains and rearing

*Acyrthosiphon pisum* aphids of the LSR1 strain were used for most experiments in the present report. LSR1 is the pea aphid strain whose genome was firstly sequenced (The International Aphid Genomics Consortium, 2010). This strain has been maintained in our lab on *Vicia fabae* seedlings for more than 8 years under long day (LD) photoperiod conditions (i.e. 16 h lights on and 8 h of darkness, or 16L:8D) at 18 °C. Strain LSR1 produces sexual females and males when reared under short day (SD) conditions (i.e. ≤ 12 h lights on and ≥ 12 h of darkness). For experimental validation of PDF sequences, additional pea aphid strains were used in addition to LSR1 (see strain details in Table S1).

### 4.2 Identification of PDF-encoding genes in aphid genomes

The PDF sequence as predicted from the *Rhodnius prolixus* genome with the surrounding convertase cleavage sites, KRNSEIINSLLGIPKVLIDAGR, was used as query in a TblastN search on the *A. pisum* NCBI Genome Reference Sequence database (Annotation Release 103, June 2019). If the putative *A. pisum* PDF sequence found would represent a functional neuropeptide, it should be conserved and expressed in other aphid species. It was therefore used as a query in a TblastN search of the Aphidomorpha transcriptome shotgun assemblies available at NCBI.

Additional Aphidomorpha PDF homologues were obtained using BlastP searches against the NCBI Protein Reference Sequence database or against the Aphid Genome Database at the BioInformatics Platform for Agroecosystem Arthropods (BIPAA, INRAE, France). We also performed tBlastn searches against whole aphid genome or transcriptome sequences at NCBI. In this later case, sequences were translated before their inclusion in the alignment (Table S2 provides Accession numbers and details of these searches). Alignment of aphid sequences (including the predicted *A. pisum* sequence) was done using ClustalX 2.0 [84]. Phylogenetic reconstructions and calculations of number of amino acid differences between aphid PDF sequences were conducted using MEGA version 11 [85].

### 4.3 Experimental validation of the pea aphid PDF-encoding gene

To experimentally validate gene models, total RNA was extracted from groups of 4-5 aphids of the above-described strains using TRI Reagent® (T9424, Sigma-Aldrich, USA) and Direct-Zol RNA extraction Kit (R2052, Zymo Research, USA) following supplier’s recommendations. RNA was quantified by spectrophotometry using the NanoDrop ND-1000 (Nanodrop Technologies, Inc., Wilmington, DE, USA) and stored at −80°C until use. 1-5 μg total RNA was used for cDNA synthesis using the NZY First-strand cDNA Synthesis Kit (ref. MB40001, NZYTech, Portugal), primed with a mix of oligo (dT)18 to enrich the samples with cDNA from mature mRNAs and hexamers. Primers to PCR amplify the predicted aphid PDF gene transcript on cDNA were designed based on 5’ and 3’ UTRs of the predicted transcript model. Forward primer was T1-F1 (TCATAATCACCGAAGACAGCAAG) and R0 (CAGTTTGTATGTGCGTTACCTACG) was the reverse primer (see Fig.1). Amplified products were directly sequenced using PCR primers after purification through 4M ammonium acetate-ethanol precipitation. Direct sequencing was done using the ABI Prism BigDye® terminator v3.1 Cycle Sequencing kit (Applied Biosystems) in an ABI3730XL sequencer. Chromatogram handling and processing was performed using the STADEN package [86]. To resolve allelic combinations present at two observed polymorphic positions, we proceeded to clone the amplified fragments for three of the strains sequenced (strains LSR1, SUT and BOL, see Table S1). For cloning the amplified fragments we used the NZY-A PCR cloning kit and NZYStar Competent Cells (refs. MB053 and MB00501 NZYTech, Portugal). Inserts contained in recombinant plasmids were sequenced as described above using plasmid based primers T7 (TAATACGACTCACTATAGGG) and M13 (GTTTTCCCAGTCACGACGT).

### 4.4 Quantification of pdf gene expression by RT-qPCR

Excised aphid heads from aphids of the LSR1 strain were used to compare the expression of the PDF gene under two photoperiodic conditions and at four different timepoints along the day. Synchronized adult aphids reared under long day (LD) and under short day (SD) conditions were sampled the following day after their final moult at ZTs 4, 11, 16 and 23. Aphid samples were kept at −80° C until RNA extraction. LD aphids had been reared under a 16L:8D regime while SD aphids were under 10L:14D conditions. For SD aphids the G1 generation was used [63]. Three groups of 5 aphids were used as replicates for each condition. Total RNA was extracted from heads from the frozen aphid samples and quantified as described above. For cDNA synthesis we used the Superscript III kit (Invitrogen) on 1 µg (c.a.) of total RNA primed with oligo (dT)18 and random hexamers following supplier’s recommendations.

RT-qPCR was performed using the AriaMx Real-Time PCR System (Agilent) and the SYBR qPCR Master Mix (ref. HY-K0501, Med Chem Express, Sweden). Primers used for RT-qPCR were QF4 (ATCCGTTGCGTACTACCTATTG) and QR4 (CATCTTCCACGTGTCTCTTACC). For each sample, three technical replicates were done. The RpL7 gene was used as an endogenous control of constitutive expression [87]. Efficiency of the PDF and of the RpL7 primer pairs were 103,7 and 90,8%, respectively. Relative expression for each sample was calculated using the ΔΔCt (threshold cycle) method [88]. All relative expression values were normalized to an inter-run calibrator sample (IRC) consisting of a cDNA synthesized from a mix of total aphid RNAs obtained from whole insects at different developmental stages. Two-way ANOVA was used to analyse the effects of photoperiod, and ZT on gene expression with SPSS Statistics 28.0 software (IBM Corp., 2021).

### 4.5 Aphid PDF antiserum and specificity assay

A polyclonal antiserum against a synthetic peptide (CSLYVPDDNFVIEEQNAPIAT) corresponding to a region of the *A. pisum* PDF Associated Peptide (Fig. 1) was raised in guinea pigs by Moravian-Biotechnology Ltd. (Brno; Czech Republic). We opted to make an antiserum to a part of the PDF precursor rather than the neuropeptide itself as it allowed for a longer peptide sequence as antigen thereby potentially increasing our chances of obtaining a good antiserum. The synthetic peptide was custom synthetized by Shanghai RoyoBiotech Co.,Ltd (Shanghai, China, 201200). After immunisation of a guinea pig and animal bleeding antisera was obtained by affinity purification column using the synthetic peptide used as immunogen.

### 4.6 Brain dissections and immunohistochemistry

Adult aphids were fixed in 4% paraformaldehyde in PBST (phosphate buffered saline PBS containing 0.5% Triton-X100) for 4 hours at room temperature (RT). They were washed 3 x 10 min in PBS (phosphate buffered saline) and then the brains were dissected in PBS. Brains were incubated in NGS solution (5% normal goat serum in PBST) for 2h at RT or over-night at 4°C.

For the immunostaining against PDF and the co-immunostainings against PDF and ILP4 or PDF and CRY, we applied the following protocol. Brains were incubated in the primary antibody solution (NGS 5%, NaN_3_ 0.02%, and primary antibodies in PBST) for two days. The following antibody dilutions were used: PDF 1:1000, PDF-CRY 1:5000 - 1:1000 respectively, and PDF-ILP4 1:5000 - 1:5000 respectively. Brains were then washed 6 x 10 min in PBST and incubated in secondary antibody solution (5% normal goat serum in PBST, Alexa Fluor 488 or 633 goat anti-guinea pig 1:200; Alexa Fluor 488 goat anti-rat 1:200; Alexa Fluor 555 or 633 goat anti-rabbit 1:200 (Thermo Scientific)) for 4h at RT, then washed 4 x 10min in PBST and 1 x 10min in PBS. Brains were then put on specimen slides and embedded in Vectashield Antifade mounting medium (Vector Laboratories, Burlingame, CA). Slides were stored at 4°C until scanning. For the co-immunostaining against PER and PDF, we applied the primary antibodies sequentially, because the PDF staining was very strong and appeared to interfere with the PER staining. First, we incubated the brains for two days at 4°C in PER primary antibody solution (NGS 5%, NaN_3_ 0.02%, PER 1:2000 in PBST) and followed the same procedure described above until the application of the secondary antibody solution (5% normal goat serum in PBST, Alexa Fluor 488 anti-guinea pig 1:200 (Thermo Scientific)). Subsequently, we washed the brains 6 x 10 min with PBST and incubated them for one day at RT with the PDF primary antibody solution (NGS 5%, NaN_3_ 0.02%, PDF 1:5000 in PBST). Then, brains were washed 6 x 10 min in PBST, incubated in secondary antibody solution for 4 hours at RT (5% normal goat serum in PBST, Alexa Fluor 633 anti-guinea pig 1:200 (Thermo Scientific)), and finally washed 4 x 10min in PBST and 1 x 10min in PBS. Brains were then put on specimen slides and embedded in Vectashield Antifade mounting medium (Vector Laboratories, Burlingame, CA). Slides were stored at 4°C until scanning. Table S3 provides details on all the antibodies used.

### 4.7 Microscopy and imaging

All the immunostainings besides the PDF-PER double-labelling were visualized with a Leica TCS SPE confocal microscope (Leica, Wetzlar, Germany). We used a 20-fold or 40-fold glycerol immersion objective (ACS APO Leica Microsystem, Wetzlar, Germany) and obtained stacks of 2 μm and 1,048,576 pixels.

For the PDF-PER double-labelling, we used a Leica CLSM SP8 (Leica Microsystems, Wetzlar, Germany). We used a 20-fold glycerol immersion objective (HC PL APO, Leica Microsystem, Wetzlar, Germany) and similarly to before, we obtained stacks of 2 μm and 1,048,576 pixels.

The confocal stacks were analyzed with Fiji ImageJ [89]. Only contrast, brightness, background correction, and color scheme adjustments were applied to the confocal images.

### 4.8 Quantification of PDF

For PDF quantification in the *pars lateralis* terminals, samples were processed in exactly the same way during the staining protocol and were scanned with identical laser settings. The quantifications were conducted in ImageJ (Fiji) and we used three different methods. First, we performed a 3D reconstruction of the PDF terminals with the Simple Neurite Tracer Plug-in (Fiji ImageJ) and defined the region of interest by creating a binary mask of the reconstruction. This was then overlapped to the maximum projection (Command: Z-projection → sum Slices) of the PDF confocal stacks and finally the average value of the pixel intensity inside the mask was calculated. Second, we used the method described in [54], that considers not only the intensity of staining in the terminals but additionally the area, which the terminals cover. For each brain, we compiled maximum projections (encompassing 10 to 15 confocal stacks), which contained the PDF terminals in the *pars lateralis* of both brain hemispheres, and subsequently removed all staining outside of them. Images were then cut to 100 000 pixels (500 pixels wide and 200 pixels high; see Figs. 4, 6) taking care that the entire PDF terminals were in the image. All resulting images were therefore of the exact same size and contained only the PDF terminals in the *pars lateralis*. We then set the background of each image to zero and measured the mean total intensity of the whole image, which reflected the extension and intensity of the dorsal projection terminals. These manipulations were done without knowing the ZT or the photoperiod at which the samples were taken to avoid any subjective influence of the investigator. We quantified at least 14 brains for each time point.

Third, we measured the length of the terminals on the compiled maximum projections (see Figs. 4, 6) by tracing a line spanning through the terminals and measuring its length (Command: Analyse → Measure) in ImageJ. For most brains, we measured the length of both PDF terminals and then calculated a mean length out of the two values. However, in few cases it was difficult to measure the terminal in one brain hemisphere since it was strongly curved. In such cases, we took only the measurement of one brain hemisphere for calculating the mean of all brains.

Two-sample t-test or Kruskal Wallis test were used to test for significant differences in PDF intensity and terminal length. The statistic tests were performed in R version 4.2.2. [90]

## Data accessibility

Sequences obtained in this work have been deposited in GenBank with accession numbers indicated in Table S1

## Acknowledgements

We thank the facilities at the Genomics, and Greenhouse services at the SCSIE (Universitat de València) for sequencing and for growing *Vicia faba* plants, respectively. We are grateful to Jochen Krauss and Ingolf Steffan-Dewenter (Animal Ecology and Tropical Biology) for the use of incubators for aphid rearing at the University of Würzburg.

