## Supplementary Table 1 3 Figure 1 for "Pigment-Dispersing Factor is present in circadian clock neurons of pea aphids and may mediate photoperiodic signalling to insulin-producing cells": Supplementary Figure 1.pdf

*** Corresponding authors:**

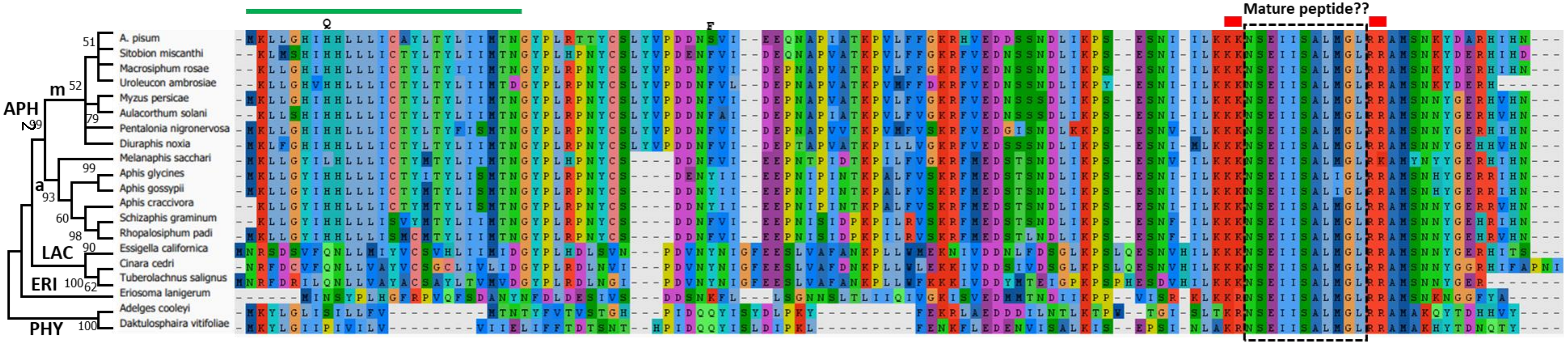


**Figure S1. Alignment of putative PDF propeptide sequences found in different aphid species.** On the left side, a cladogram inferred from the aligned sequences is depicted. The *A. pisum* predicted sequence is at the top position in the alignment. The two positions found polymorphic in *A. pisum* are indicated. The green line marks the predicted signal peptide in *A. pisum*. The red thick lines indicate the predicted convertase cleavage sites in A. pisum. 11 aminoacids highly conserved in all aphids are included in a dotted black box and denoted as Mature peptide ?? (see text for details). Some groupings are indicated. APH, subfamily Aphidinae; LAC, subfamily Lachninae; ERI, subfamily Eriosomatinae; PHY, superfamily Phylloxeroidea (containing oviparous Adelgidae and Phylloxeridae); m, tribe Macrosiphini; a; tribe Aphidini. Accession numbers of the aligned sequences are indicated in Table S2.
