## Supplementary Table 1 3 Figure 1 for "Pigment-Dispersing Factor is present in circadian clock neurons of pea aphids and may mediate photoperiodic signalling to insulin-producing cells": Supplementary Tables 1-3.pdf

*** Corresponding authors:**

**Table S1:** Information on different pea aphid strains used in the present work including the aminoacids present at the two polymorphic positions (Pos. 8 and Pos. 43, see Fig. 1) in the PDF propeptide. * Indicates whether the aphids are holocyclic (Holo, they have a sexual phase) or anholocyclic (Anholo, they lack a sexual phase). ** Location, indicates where the original founding aphids were first collected. T1 acc. nº, indicates accession numbers for transcript 1.

| Strain | Cycle* | Location** | Colour | Pos. 8 | Pos. 43 | T1 acc. nº |
|---|---|---|---|---|---|---|
| LSR1 | Holo | Ithaca (USA) | Red | H/Q | F/S | pending |
| Bol | Holo | York (UK) | Green | Q | F | pending |
| RAS | Holo | York (UK) | Green | H/Q | F/S | pending |
| SUT | Holo | York (UK) | Green | H/Q | F/S | pending |
| GR | Anholo | Gallur (Spain) | Red | H | S | pending |
| Peña | Anholo | Peñaroya (Spain) | Green | H/Q | F/S | pending |
| YR2 | Holo | York (UK) | Red | H/Q? | F/S | pending |

**Table S2:** accession numbers and details of PDF sequences from different aphid species used in Fig. S1

| Aphid species | Database | accession | Method | Comment |
|---|---|---|---|---|
| *Adelges cooleyi* | NCBI | XP_050429540 | BlastP | |
| *Aphis craccivora* | NCBI | VUJU01002760* | tBlastn | whole genome shotgun |
| *Aphis glycines* | BIPAA | AG007432-PA | BlastP | |
| *Aphis gossypii* | NCBI | XP_027837487 | BlastP | |
| *Aulacorthum solani* | NCBI | PVMI01058046* | tBlastn | whole genome shotgun |
| *Cinara cedri* | NCBI | CABPRJ010000953* | tBlastn | whole genome shotgun |
| *Daktulosfaira vitifoliae* | NCBI | GDEB01008642.1 | tBlastn | Transcriptome |
| *Diuraphis noxia* | NCBI | XP_015364753 | BlastP | |
| *Eriosoma lanigerum* | BIPAA | jg20706.p1 | BlastP | |
| *Essigella californica* | NCBI | GAZF02015936** | tBlastn | Transcriptome Shotgun Assembly |
| *Macrosiphum rosae* | NCBI | WHPZ01537923* | tBlastn | whole genome shotgun |
| *Melanaphis sacchari* | NCBI | XP_025209065 | BlastP | |
| *Myzus persicae* | NCBI | XP_022174727 | BlastP | |
| *Pentalonia nigronervosa* | BIPAA | g557.p1 | BlastP | |
| *Rhopalosiphum padi* | NCBI | XP_026814266 | BlastP | |
| *Schizaphis graminum* | NCBI | VSBP01000111 * | tBlastn | whole genome shotgun |
| *Sitobion miscanthi* | NCBI | SSSL01000001* | tBlastn | whole genome shotgun |
| *Tuberolachnus salignus* | NCBI | GCYY01004835** | tBlastn | Transcriptome Shotgun Assembly |
| *Uroleucon ambrosiae* | NCBI | GCZC01002829** | tBlastn | Transcriptome Shotgun Assembly |

* Genomic sequence
** Transcribed RNA

**Table S3.** Details on antibodies used in the present report

| Antibody | Host | Reference | RRID |
|---|---|---|---|
| *anti*-PDF | Guinea pig | present article | |
| *anti*-PER | Rat | Colizzi et al, 2021 | AB_2314242 |
| *anti*-CRY | Rabbit | Yoshii et al, 2008 | AB_2920831 |
| *anti*-ILP4 | Rabbit | Cuti et al, 2021 | |
